# A plasma membrane nanoplatform ensures signal specificity during osmotic signaling in plants

**DOI:** 10.1101/2020.06.14.145961

**Authors:** M. Smokvarska, C. Francis, M.P. Platre, J.B. Fiche, C. Alcon, X. Dumont, P. Nacry, V. Bayle, M. Nollmann, C. Maurel, Y. Jaillais, A. Martiniere

## Abstract

In the course of their growth and development plants have to constantly perceive and react to their environment. This is achieved in cells, by the coordination of complex combinatorial signaling networks. However, how signal integration and specificity are achieved in this context is unknown. With a focus on the hyperosmotic stimulus, we use live super-resolution light imaging methods to demonstrate that a Rho GTPase, Rho-of-Plant 6 (ROP6), forms stimuli-dependant nanodomains within the PM. These nanodomains are necessary and sufficient to transduce production of reactive oxygen species (ROS),that act as secondary messengers and trigger several plant adaptive responses to osmotic constraints. Furthermore, ROP6 activation triggers the nanoclustering of two NADPH oxidases that subsequently generates ROS. ROP6 nanoclustering is also needed for cell surface auxin signaling, but short-time auxin treatment does not induce ROS accumulation. We show that auxin-induced ROP6 nanodomains, unlike osmotically-driven ROP6 clusters, do not recruit the NADPH oxidase, RBOHD. Together, our results suggest that Rho GTPase nano-partitioning at the PM ensure signal specificity downstream of independent stimuli.

## INTRODUCTION

Biological membranes can be seen as a patchwork where lipids and proteins are grouped in a juxtaposition of domains of various shapes and sizes. Paradoxically, membranes are also a fluid-structure allowing lateral diffusion of its constituents by thermal agitation. This property of membranes is central as it participates in the dynamic partitioning of proteins and lipids between different plasma membrane (PM) domains and consequently regulates cell-surface signaling processes [1]. In plants, the vast majority of PM proteins observed with improved fluorescent microscopy technics was described to be organized in nanodomains of long dwell time (several minutes). It is especially the case of REMORIN3.1 (REM3.1), PLASMA MEMBRANE INTRINSIC PROTEIN2;1 (PIP2;1), PIN-FORMED2 (PIN2), AMMONIUM TRANSPORTER3.1 (AMT3.1), BRASSINOSTEROID INSENSITIVE1 (BRI1), RESPIRATORY BURST OXIDASE HOMOLOG PROTEIN D (RBOHD), FLAGELLIN SENSING2 (FLS2) and NITRATE TRANSPORTER1.1 (NRT1) [2–8]. Nevertheless, the functional relevance of this particular membrane organization remains poorly understood and its role in cell signaling just begins to be explored.

Among other signals, plant cells respond to changes in water availability generated by osmotic constraints. Despite tremendous effort in the last decades, the molecular mechanisms that allow plant cells to perceive and induce early signaling events in response to osmotic stress just begin to be understood [9,10]. One of the first cellular responses is an accumulation of reactive oxygen species (ROS) [11] in cells, which act as secondary messengers, regulating cell endocytosis but also root water conductivity and intracellular accumulation of osmotica (e.g. proline) [12,13]. Two processes are under action to generate ROS during osmotic signalling. One is non-enzymatic and requires reduction of apoplastic iron. The second is mediated by the PM-localized NADPH oxidases, RBOHD and F [11]. RBOHs catalyze the production of superoxide free radicals by transferring one electron to oxygen from the cytoplasmic NADPH. Even if the mechanism that drives ROS production is now better understood, it is still unclear how it is triggered by a change in osmolarity.

The Rho of plant (ROP), belonging to the super clade Ras/Rho GTPase, have a key role in cell surface signaling events including response to hormones such as auxin or ABA, but also during biotic stimulation [14]. In some cases, they also appear to regulate ROS accumulation, like in tip growing cells or in response to chitin elicitation [15,16]. ROPs are functioning as molecular switches due to a change in conformation between an active GTP-bound form and an inactive GDP-bound form. But, ROP function is also tightly associated with its lipid environment. For instance, the rice type-II ROP OsRAC1interacts with OsRBOHB in the presence of specific sphingolipids containing 2-hydroxy fatty acids [17]. Besides, the role of charged lipids was recently exemplified in a work on a type-I ROP from *Arabidopsis thaliana*. In this study, the anionic lipid phosphatidylserine (PS) was shown to interact directly with ROP6 C-terminal hypervariable domain, to determine ROP6 organization at the PM and to quantitatively control plant response to the phytohormone auxin [18]. Therefore, *ROP* gene family constitutes a good candidate to regulate osmotic signaling.

Here, we show that ROP6 is a master regulator of osmotically-induced ROS accumulation and participates in a set of plant responses to this signal. Using super-resolution microscopy, we found that ROP6 co-exists in the same cell in different states and that osmotic stimulation induces ROP6 nanodomain formation. These nanodomains are needed for a correct ROS accumulation in cells and their composition differs when triggered by other stimulus, suggesting that ROP6 nanodomains may encode for signal specificity.

## RESULTS

### ROP6 is necessary for osmotically induced ROS accumulation and participates in plant responses to osmotic signal

To investigate the potential role of ROPs in osmotic signaling, we used medium or high sorbitol concentration (ψ_medium_= -0.26 MPa and ψ_high_= -0.75 MPa, respectively), and challenged *rop* loss-of-function mutant lines corresponding to the three isoforms that are highly expressed in roots (SF1 A). ROS accumulation in cells, as revealed by DHE dye, was used as a fast readout for activation of osmotic signalling [11]. Compared to WT, *rop6.2* seedling, but not *rop2.1* nor *rop4.1*, shows impaired ROS accumulation, (Figure 1A-C, FigS1 B). No additive effect was detected in *rop2.1xrop6.2* or *rop2.1xrop6.2*xROP4RNAi (FigS1 B). A defect in ROS accumulation was fully complemented by a transgene containing mCitrine-tagged *ROP6* genomic DNA driven by its promoter (*rop6.*2xmCit-ROP6, Figure 1C).

**Figure 1:**
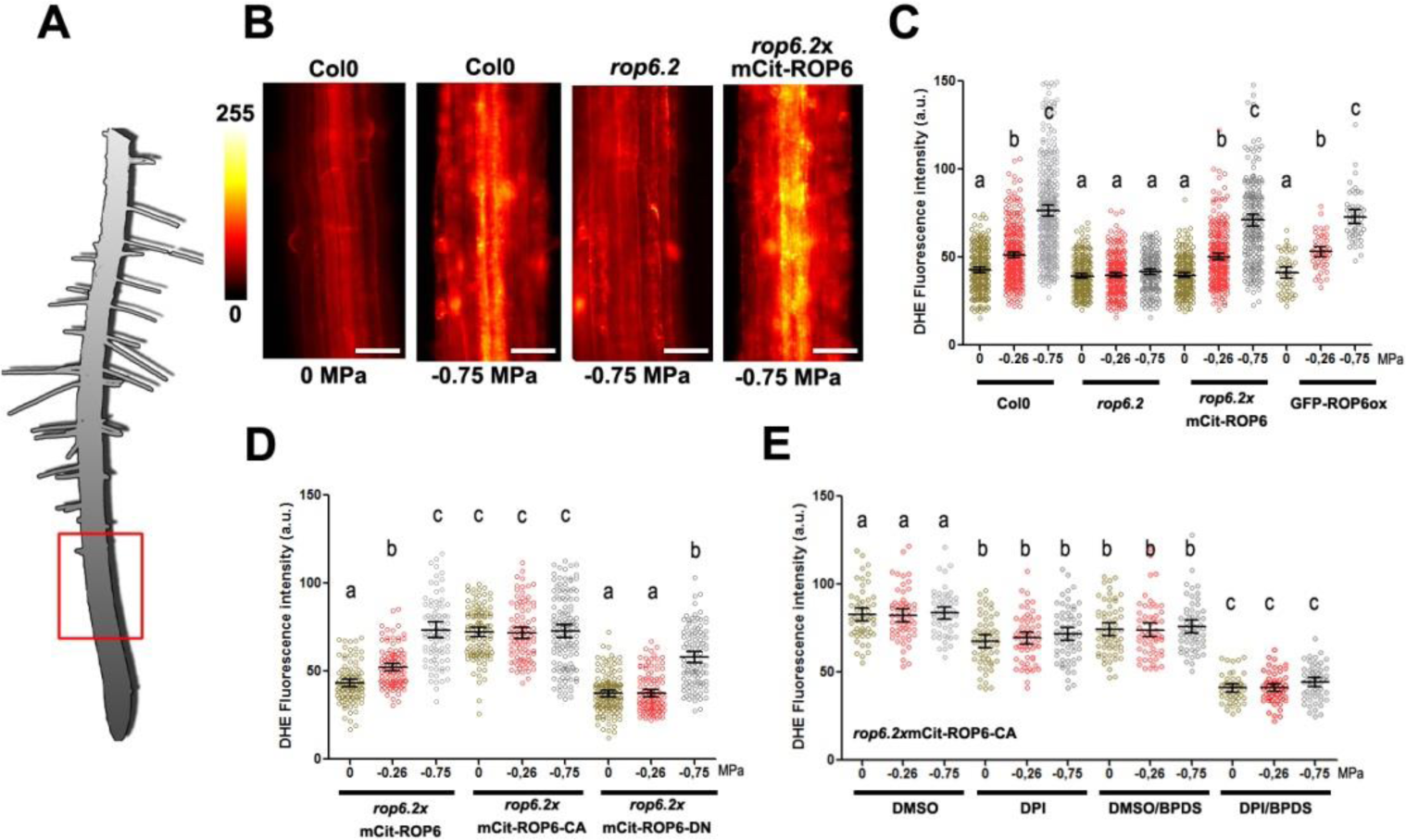
ROP6 activation is necessary and sufficient to trigger osmotically driven ROS accumulation in Arabidopsis root cells. (A) Drawing of Arabidopsis plantlets, where the red square highlights the part of root understudy. (B) Dihydroethidium (DHE) stained root cell of Col0, *rop6.2* and *rop6.2*xmCit-ROP6 in control condition (0 MPa) or after 15 min of -0.75 MPa treatment. (C-D) DHE fluorescence quantification after 15 min treatment with 0, -0.26 or -0.75 MPa solution in different genetic material: Col(0), *rop6.2*, ROP6 over expresser line (GFP-ROP6ox) and *rop6.2* lines expressing under ROP6 endogenous promotor (mCit-ROP6 (*rop6.2*xmCit-ROP6), the constitutive active ROP6 (*rop6.2*xmCit-ROP6-CA) or the dominant negative (*rop6.2*xmCit-ROP6-DN). (E) ROS quantification (DHE fluorescence) in root cell expressing the constitutively active ROP6 (mCit-ROP6-CA) in control or after mild or high osmotic stimulus (respectively 0, -0.26 and -0.75 MPa) supplemented or not with ROS enzyme inhibitor. DPI was used for inhibition of NADPH oxidase activity and BPDS to inhibit ROS produce from ferric iron. Error bars correspond to a confidence interval at 95%. ANOVA followed by Tukey test, letters indicate significant differences among means (p-value<0.001). N=4-6 independent biological replica. Scale bar 20 µm.

Because, in roots, ROS accumulation has been tightly associated with the deposition of the secondary wall, especially lignin [19]. We wondered if osmotic constrain could enhance cell lignification. Roots exposed to -0.75 MPa for 24 hours have a strong autofluorescence signal compared to control situation, and when stained with phloroglucinol that reveal lignin specifically, typical cherry-red staining was observed [20] (FigS2 A and B). We tested if the osmotically-enhanced lignin deposition is indeed associated with ROS accumulation. Plants invalidated for the two highly expressed NADPH oxidases (*RBOHD* and *F)*, showed a reduced autofluorescence after exposure to - 0.75 MPa and control plants exposed to 1mM H_2_0_2_ for an hour revealed a strong fluorescent signal, showing a connection between osmotically-induced lignin deposition and ROS production (FigS2 C). This response was partially regulated by ROP6, as *rop6-2* plants show dimmer root autofluorescense signal after -0.75 MPa treatments than in control plants either Col0 or *rop6.2*xmCit-ROP6 (FigS2 C-D).

Interestingly, after 48 hours on -0.75 MPa plate, root tip cells displayed local isotropic cellular growth (FigS2E). This change in cell polarity has been suggest to reflect the acclimation process of the root facing hyperosmotic condition, as it was described for salt or drought responses [21,22]. Because, ROPs are known to regulate cell polar growth of pavement cells, pollen tube and root hairs [14], we wondered if *ROP6* may participate in the osmotically induced cell isotropic growth. *rop6.2* shows a significantly smaller circularity index that wild type or complemented lines on treated plate (−0.75 MPa), whereas no difference between genotypes was found in control conditions (FigS2 F and G).

Because, ROP6 seems to participate in multiple phenotypes associated to plant acclimation to osmotic constrain, we wondered if *ROP6* can also participates in the changes of root growth and development. Whereas indistinguishable when 5DAG plants were transplanted in control conditions, *rop6.2* plants grew slightly faster than *rop6.2*xmCit-ROP6 in -0.75 MPa plate (rate constant^*rop6.2*xmCit-ROP6^ =0.011^-h^+/-0.0005^-h^, rate constant^*rop6.2*^=0.009^-h^ +/-0.0008^-h^, T-test p-value=0.02, SF2H-K). Indeed, plants have longer primary and lateral roots in loss-of-function *rop6.2* mutant in this stress condition, while no significant effect was observed for lateral root density (FigS2 L-M). Interestingly, *ROP6* expression pattern fits a potential role in root growth, as mCit-ROP6 fluorescence is mostly present in the root meristem and elongation zone and in lateral root primordia (FigS3 A-E). As a whole, ROP6 appears to be necessary for osmotically-induced ROS accumulation, but also participate in plant adaptations to hyperosmotic treatment (FigS2 D, G and K).

### ROP6 activation, but not protein quantity, is rate-limiting to trigger osmotic signaling

Next, we tested if *ROP6* is sufficient to trigger osmotic signaling. Although GFP-ROP6ox overexpressing lines accumulate high amounts of ROP6 proteins, no enhancement of osmotically-induced ROS was observed in control condition or after treatment, suggesting that ROP activation rather than protein quantity might be a limiting factor (FigS4 A and Fig1 C). To test this hypothesis, we used *rop6.2* lines expressing, under a native ROP6 promotor, constitutive active GTP-lock (ROP6-CA) or constitutive inactive GDP-lock ROP6 dominant negative version (ROP6-DN) in fusion with mCitrine. Plants expressing mCit-ROP6-CA showed a constitutively high ROS accumulation, independent of the stimulus (Fig1 D). Oppositely, in mCit-ROP6-DN plants, ROS induction was attenuated after exposure to -0.75 MPa and totally suppressed after -0.26 MPa treatments, compared to a control situation (Fig 1 D). These results showed that ROP6 is necessary and its activation sufficient to trigger ROS production. Then, we addressed if ROP6 activation could act upstream of ROS producing enzymes. Therefore, rop6.2xmCit-ROP6-CA line, that has constitutively high ROS, was treated alone or in combination with specific inhibitors for each of the two ROS pathways activated by the osmotic stimulus [11]. Diphenyleneiodonium (DPI) was used to inhibit NADPH oxidase activity and bathophenanthrolinedisulfonic acid (BPDS) to block ROS mediated by ferric iron[11]. In co-treatment, ROS generated by mCit-ROP6-CA is diminished drastically, suggesting that mCit-ROP6-CA is acting upstream of ROS production machinery (Fig1 E). Next, we determined if ROP6 activation is associated with a change in its subcellular localization, as described for many small GTPases[23]. A sharp fluorescent signal was observed delimiting root cells expressing mCit-ROP6, which overlaid with the FM4-64 PM dye (FigS3 F). No difference in fluorescence intensity was detected at the PM between wild type and GTP or GDP lock ROP6 (FigS4 B), suggesting that ROP6 activation has a minor role in ROP6 PM targeting.

### Two populations of ROP6 molecules co-exist within the plasma membrane and vary in frequency minutes after osmotic treatment

It was recently shown that ROP6 organization at the PM is critical for auxin signaling^16^. We thus addressed whether ROP6 lateral segregation at the PM could contribute to osmotic signaling. Total internal reflexion fluorescent microscopy (TIRFM) showed that GFP-ROP6 has a uniform localization within the PM in control conditions, while in -0.26 MPa and even more in -0.75 MPa treated cells, GFP-ROP6 appeared in diffraction-limited spots at the cell surface (Fig2 A). This suggests that ROP6 clustered in response to osmoticum treatment in a dose-dependent manner (Fig2 B). Kymogram analysis showed straight lines for up to 50 seconds, suggesting that GFP-ROP6 clusters are stable within the PM during this period (Fig2 C). The average GFP-ROP6 spot size is close to the limit of diffraction (radius = 235 +/- 60.57 nm), therefore we next used sptPALM, a super-resolution imaging technic, recently developed on plant sample [11,18,24]. Upon stochastic photoswitching on live roots expressing mEOS2-ROP6, sub-diffraction spots are appearing with blinking behaviour and small life span (< 0.5sec), as expected from single molecule behaviour (Supp Video1 and FigS5 A and B). By retrieving the displacement of each ROP6 single molecule along with the videos, two behaviours can be observed in control condition (Fig2 D, highly diffusible molecules in orange and lowly diffusible molecules in blue, Fig2 E). Distribution of instantaneous diffusion coefficient of ROP6 single molecules, extrapolated from mean square displacement plots, is bimodal (Fig2 F, green curve Fig2 G). This result shows that diffusible (D_diff_=0.05+/-0.007µm^2^.s^-1^) and relatively immobile (D_imm_=0.002+/-0.0007µm^2^.s^-1^) mEOS2-ROP6 molecules coexist within the PM of a single cell. Minutes after -0.75 MPa treatments, the frequency of immobile mEOS2-ROP6 doubles (Fig2 G, H and I). Clustering analysis on live PALM images, using Vonoroï tessellation [25] (Fig2 J), showed that the occurrence of molecules with high local density increases after -0.75 MPa treatment (Fig2 K and L, Log_local density_ >3). Nevertheless, at this stage it was not possible to distinguish between three different cases: (i) the sizes of nanodomains are increasing after treatment, (ii) cells have the same number of nanodomains between control and treatment but with more ROP6 molecules in it or (iii) more nanodomains are formed in response to -0.75 MPa, with a similar amount of ROP6 protein. To distinguish between these possibilities, segmented images were generated based on detection local density, where only ROP6 molecules with a local density higher than Log_local density_>3 were investigated (FigS5 C-E). Whereas no effect on domain size, nor percentage of mEOS-ROP6 molecules per nanodomains was found, the density of nanodomains per µm^2^ of PM doubles after osmotic treatment (Fig2 M-O). Together our results suggest that in response to osmotic stimulation, ROP6 molecules are clustering in nanometer-sized domains (i.e. nanodomain), with a relatively fixed size and constant number of ROP6 molecules, and in which ROP6 barely diffuses.

**Figure 2:**
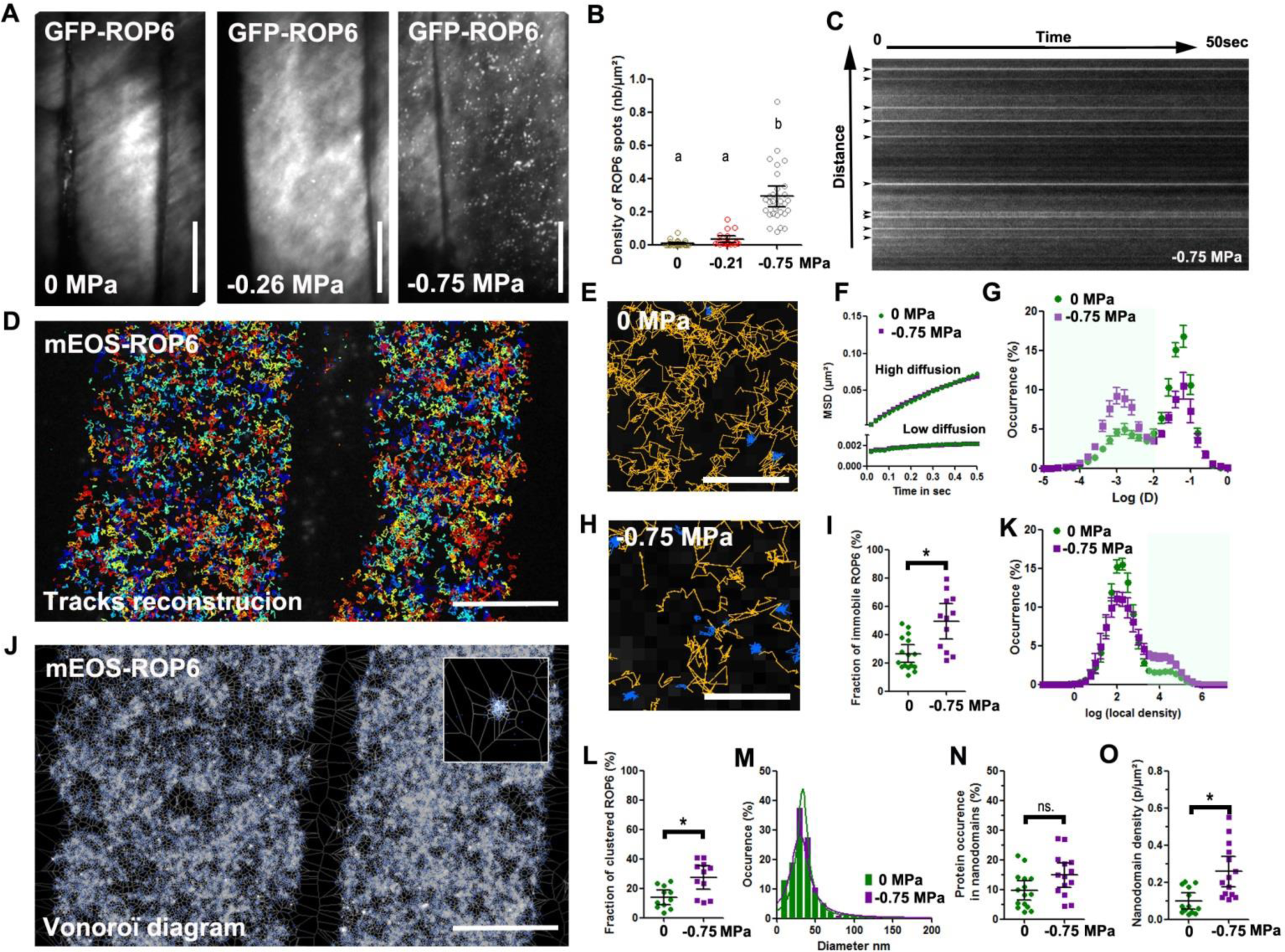
Osmotic stimulus triggers ROP6 molecular nanoclustering at the PM. (A) TIRFM micrograph of GFP-ROP6 expressing cells after 2 minutes incubation with solutions at either 0 MPa, -0.26 or -0.75 MPa. (B) Quantification of ROP6 spots density. (C) Kymograph image of GFP-ROP6 spots from cells exposed to 0.75 MPa. Spots at initial time point are labelled with arrows. (D) Image reconstruction of around 5 000 single mEOS2-ROP6 molecule trajectories in two control cells. (E) Close-up view of cell expressing mEOS2-ROP6, where trajectories with high or low diffusion instantaneous coefficient labelled in orange or blue respectively. (F) Mean square displacement curves of the highly or lowly diffusible molecules in control (0 MPa) or treatment (−0.75 MPa). (G) Bimodal distribution of molecules instantaneous diffusion coefficients in control (0 MPa, green curve) and treatment (−0.75 MPa, purple curve). (H) Close up view of the PM of cell expressing mEOS2-ROP6 2 minutes after a -0.75 MPa treatment. (I) Histogram represents the percentage of molecules with an instantaneous diffusion below 0.01 um^2^.s^-1^ in control (0 MPa) or after treatment (−0.75 MPa). (J) Vonoroï tessellation of mEOS2-ROP6 molecules localization map from the exact two control cells in (D). Top right inset is a close up view, showing mEOS2-ROP6 nanodomain. (K) Distribution of molecules local density in control (0 MPa, green curve) and treatment (−0.75 MPa, purple curve). (L) Percentage of molecules with a log(local density) higher than 3. (M) Distribution of the mEOS2-ROP6 nanodomains diameter in control (0 MPa) and treatment (−0.75 MPa). (N) Relative occurrence of mEOS2-ROP6 in nanodomains in control (0 MPa) and treatment (−0.75 MPa). (O) Nanodomain density in control (0 MPa) or after 2 minutes treatment with -0.75 MPa solution. Error bars correspond to a confidence interval at 95%. For (B) an ANOVA followed by Tukey test was done, letters indicate significant differences among means (p-value<0.001). * p-value below 0.01 T-Test. N=3 independent biological replica. Scale Bar 10µm, except for E and H where it is 1 µm

### ROP6 nanodomains are necessary to trigger osmotically-induced ROS

Next, we addressed whether ROP6-containing nanodomains are involved in osmotic signaling. Because GTP-locked ROP6 (ROP6-CA) is constitutively producing ROS (Fig1 C), we quantified diffusion and local density of mEOS2-ROP6-CA molecules by sptPALM. In comparison to the wild-type protein, ROP6-CA has a higher proportion of immobile molecules and a bigger fraction of molecules with high local density, suggesting that ROP6-CA constitutively forms nanodomains (Fig3 A, B, C and D). In addition to its C-terminal prenylation, ROP6 is transitory S-acylated on cysteines 21 and 156 [16]. These modifications are required for localization in detergent-resistant membranes and cause retarded lateral diffusion of the constitutive active GTP-lock ROP6 but have no impact on ROP6 GTPase activity or PM targeting [16]. To test if ROP6 acylation is required for nanoclustering, we generated mEOS2-ROP6^C21A/C156A^ expressing plants. Using sptPALM and clustering analysis, we found that mEOS2-ROP6^C21A/C156A^ was insensitive to -0.75 MPa treatments (Fig3 E, F, G and J). Because mEOS2-ROP6^C21A/C156A^ is not forming nanodomain in response to osmotic treatment, we compared the ROS response in *rop6.2*xmCit-ROP6^C21A/C156A^ and *rop6.2*xmCit-ROP6 complemented lines 0.26 MPa or - 0.75 MPa treatments did not trigger any ROS accumulation in *rop6.2*xmCit-ROP6^C21A/C156A^. Importantly, mCit-ROP6^C21A/C156A^ expressed under the control of its own promoter localized at the PM in root cells (Fig3 J), as previously reported for 35S::GFP-ROP6^C21A/C156A^ in leaves [16]. Together, our results suggest that ROP6 nanodomain formation, rather than only ROP6 PM localization, is necessary to activate osmotic signaling in cells.

**Figure 3:**
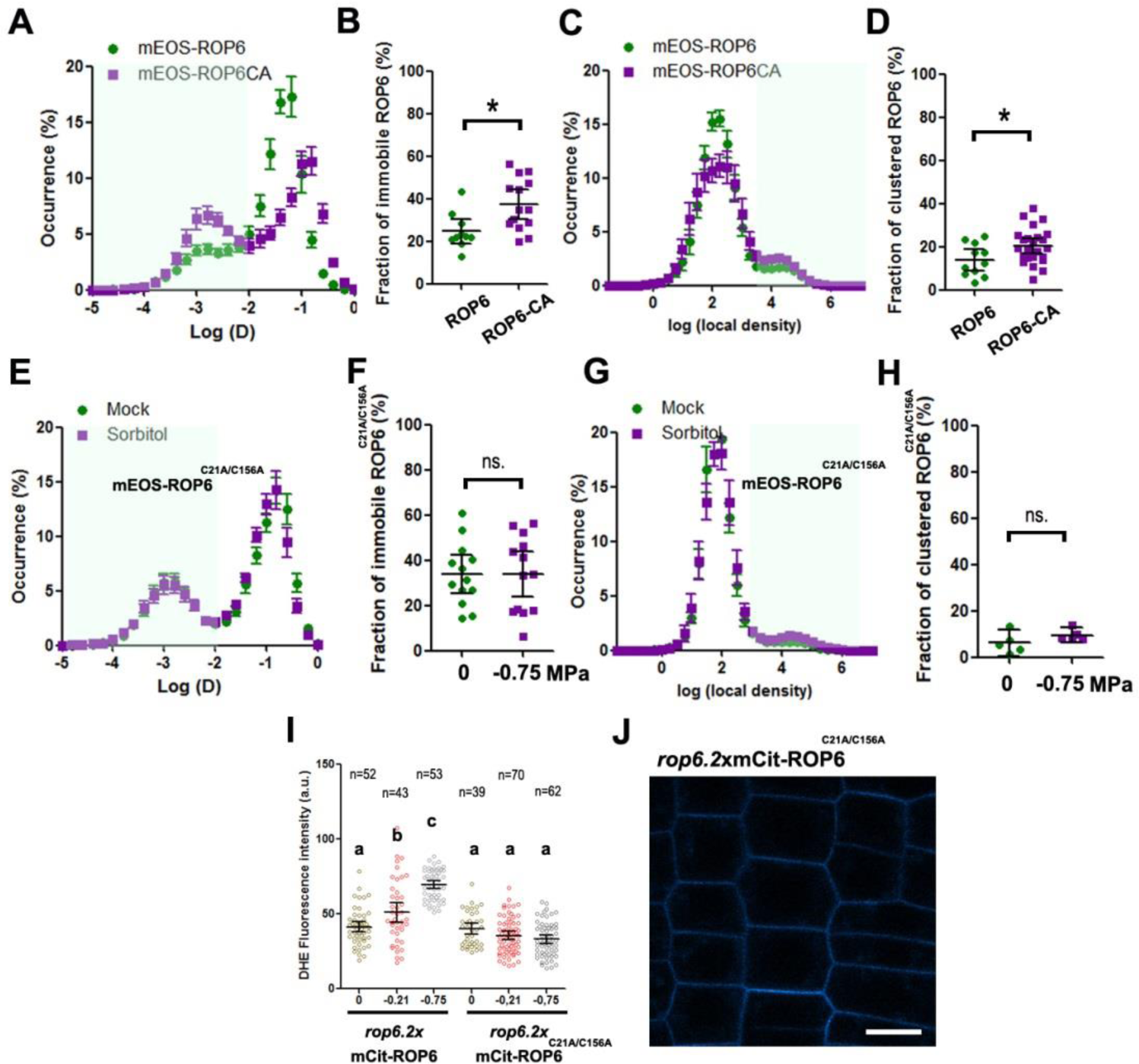
ROP6 nanoclustering is required for ROS accumulation. (A) Bimodal distribution of mEOS2-ROP6 instantaneous diffusion (green curve) or mEOS2-ROP6-CA (purple curve). (B) Histogram represents the percentage of molecules with an instantaneous diffusion below 0.01 um^2^.s^-1^ in mEOS2-ROP6 and mEOS2-ROP6-CA expressing lines. (C) Distribution of molecules local density in mEOS2-ROP6 (green curve) and mEOS2-ROP6-CA (purple curve). (D) Percentage of molecules with a log(local density) higher than 3. (E) Bimodal distribution of mEOS2-ROP6^C21A/C156A^ instantaneous diffusion coefficients in control (0 MPa, green curve) and treatment (−0.75 MPa, purple curve). (F) Histogram represents the percentage of mEOS2-ROP6^C21A/C156A^ molecules with an instantaneous diffusion below 0.01 um^2^.s^-1^ in control (0 MPa) and treatment (−0.75 MPa). (G) Distribution of mEOS2-ROP6^C21A/C156A^ molecules local density in control (0 MPa, green curve) and treatment (−0.75 MPa, purple curve). (H) Percentage of molecules with a log(local density) higher than 3. (I) Quantification of ROS accumulation by DHE staining in *rop6.2*xmCit-ROP6 or *rop6.2*xmCit-ROP6^C21A/C156A^ expressing cells after 15 min treatment with 0, -0.26 or -0.75 MPa solution. (J) Plasma membrane localization of mCit-ROP6^C21A/C156A^. Error bars correspond to a confidence interval at 95%. * p-value below 0.01 T-Test. ns. Non-significant. N=3-4 independent biological replica. Scale bar 10µm.

### Activated ROP6 interacts with RBOHD and F in PM nanodomains to generate ROS

We checked first if *ROP6, RBOHD* and *RBOHF* are co-expressed in similar Arabidopsis root cells. Transcriptional fusion for RBOHD, and translational fusion for ROP6 and RBOHF all showed signal in root epidermis (FigS6 A-C). Next, we tested if the two NADPH oxidases isoforms that are activated by osmotic signal, RBOHD and RBOHF, could interact with ROP6. FLIM experiments were performed in tobacco leave cells that transiently expressed the two putative interacting proteins tagged with GFP or mRFP. We found a significant diminution of GFP life time when GFP-RBOHD was co-expressed with RFP-ROP6-CA compared to cell expressing GFP-RBOHD and RFP-ROP6-DN or when cell expressed only the donor GFP-RBOHD (Figure 4 A and B). Similar results were observed with GFP-RBOHF, suggesting that both RBOHs interact *in planta* with the GTP-, but not the GDP-locked form of ROP6 (FigS7 A). This is in line with recent observations made by yeast two hybrids experiments, where RBOHD and ROP6-CA were shown to interact [26].

**Figure 4:**
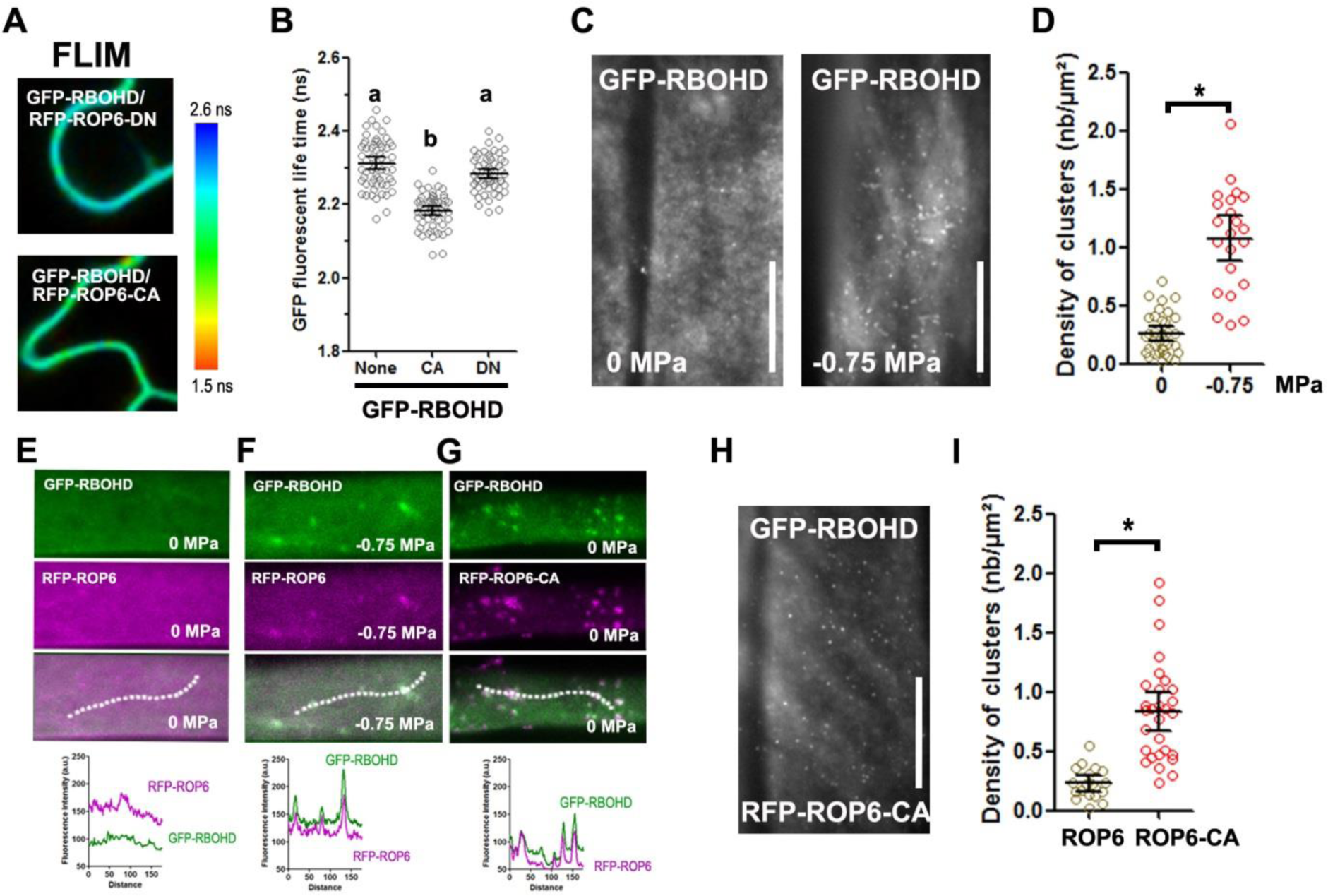
ROP6 interacts and forms nanoclusters with RBOHD at the PM. (A) GFP-RBOHD fluorescence lifetime when co-express with dominant negative (RFP-ROP6-DN) or constitutive active ROP6 (RFP-ROP6-DN) in transient expression in tobacco leaf epidermal cells and its quantification (B). (C) TIRF micrograph of cell expressing GFP-RBOHD in control or after 2 minutes treatment with -0.75 MPa solution and quantification of spots density (D). (E-G) Cell co-expressing GFP-RBOHD with RFP-ROP6 in control (E), -0.75 MPa treatment (F) or with RFP-ROP6-CA (G). Graph below represents the pixel intensity along the dotted line in each of the condition. (H) TIRFM micrograph of GFP-RBOHD signal in GFP-RBOHDxRFP-ROP6-CA plant. (I) Spot density quantification in GFP-RBOHDxRFP-ROP6 or GFP-RBOHDxRFP-ROP6-CA. Error bars correspond to a confidence interval at 95%. For (B) an ANOVA followed by Tukey test was done, letters indicate significant differences among means (p-value<0.001). * p-value below 0.01 T-Test. N=3 independent biological replica. Scale bar 10µm.

Because ROP6 and RBOHs physically interact and ROP6 forms nanodomains that are necessary for ROS accumulation, we hypothesized that RBOHs could also be organized in nanodomains in the cell PM. Arabidopsis lines overexpressing GFP-tagged RBOHD and RBOHF were generated. Under TIRF illumination, GFP-RBOHD showed a uniform localization in control condition, while 2 minutes after -0.75 MPa treatment, cells had clearly visible spots (Fig 4 C and D). By using GFP-RBOHDxRFP-ROP6 plants, we observed that ROP6 accumulated in the same structure as RBOHD after osmotic stimulation (Fig4 E and F). To analyse whether RBOH domains formation is a consequence of ROP6 activation or if an independent pathway drives it, we crossed col0 GFP-RBOHD with RFP-ROP6-CA lines. Both, RBOHD and ROP6-CA accumulated in nanodomain in the absence of osmotic stimulation and co-localize in this structure (Fig4 G). In addition, the density of RFP-RBOHD spots is much higher when the constitutively active form of ROP6 is present in cells, even in the absence of any stimulation (Fig4H and I). As *rbohF* and *rbohD* mutant plants display similar reduced ROS accumulation in response to osmotic stimulation, we tested if RBOHF would form stimuli-dependant spots in the PM, like RBOHD does [11]. Even if it has a substantial number of detectable spots in control condition, GFP-RBOHF overexpressing plant clearly showed an increased spot density minute after -0.75 MPa treatment (FigS7 B and C). This last result suggests that to some extend RBOHD and RBOHF have similar re-localisation behaviour in response to osmotic stimulation and that ROP6 activation is required to trigger this re-localization.

### Can ROP6 nanodomain formation mediate independent signaling events?

ROP6 is necessary for several plants signaling responses including to the phytohormone auxin [14,18,27,28]. The correct targeting of the auxin transport efflux carrier PIN2 is mediated by ROP6 and therefore participates in root gravitropic response [27,28]. Recently, ROP6 nanodomain formation, mediated by the anionic lipid phosphatidylserine (PS), was described in response to auxin[18]. Together with our results on osmotic signaling, this suggests that nanodomain formation is a general feature for ROP6 signaling pathways in plant. We addressed whether RBOHD clustering is also induced in response to auxin stimulation, as it happens after the induction of osmotic signaling pathway. No increase of GFP-RBOHD spots density were observed in such condition, whereas ROP6 clearly show, as expected, numerous dotted structure in the PM (Fig5 A and B). As it was previously described, roots exposed to auxin for a short time (60 min) failed to accumulate ROS, which contrasts with osmotic stimulation (Fig5 C) [29–32]. These results show that ROP6 nanoclusters formed after auxin or osmotic stimulation can differ in their constituent and consequently encode, to a certain extent, for signal specificity.

**Figure 5:**
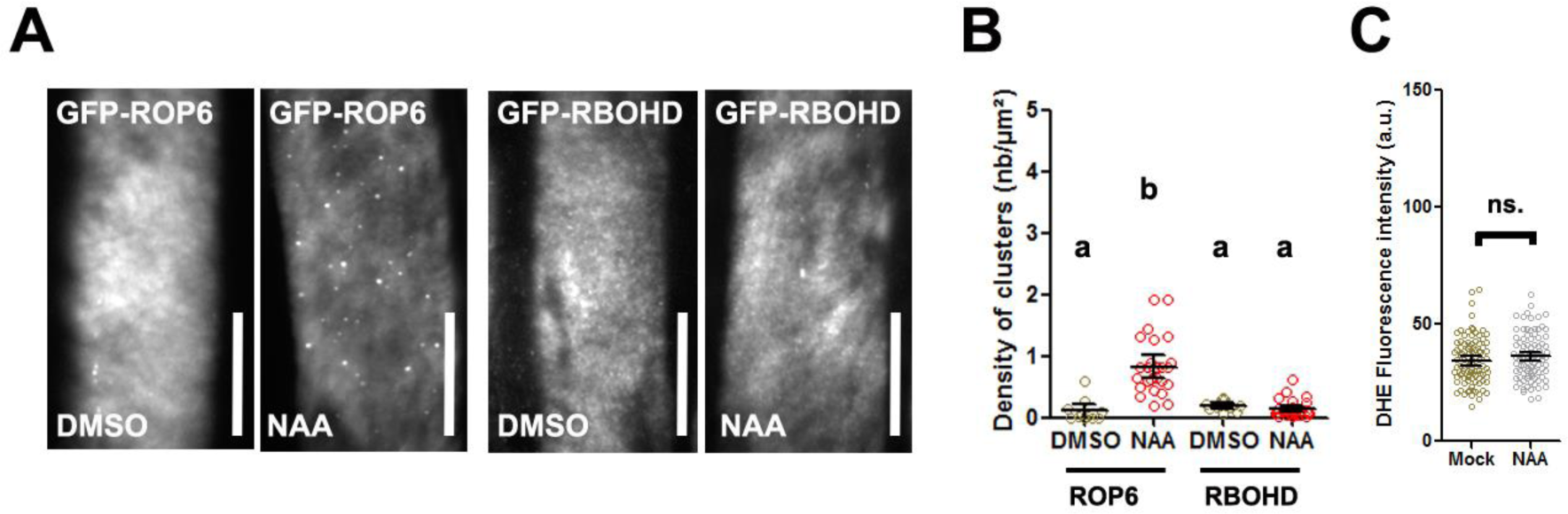
Auxin-stimulated ROP6 nanodomains are exempt from RBOHD. (A) TIRFM micrograph of cell expressing GFP-ROP6 or GFP-RBOHD in control (DMSO) or 10 μM NAA for 1 hour. (B) Spots density quantification. (C) Quantification of ROS accumulation by DHE staining in control (DMSO) or after 15 min treatment with 10 µM NAA. Error bars correspond to a confidence interval at 95%. For (B) an ANOVA followed by Tukey test was done, letters indicate significant differences among means (p-value<0.001). * p-value below 0.01 T-Test. independent biological replica. Scale bar 10µm.

## DISCUSSION

By combining genetic and super resolution live imaging, we showed that ROP6 forms osmotic specific nanodomains within the PM that are required to trigger secondary messenger in cells. The role of this specific ROP isoform is central for osmotic signaling since *rop6.2* has a totally abolished osmotically-induce ROS production. In addition, ROP2 and ROP4 which are also highly expressed in roots are dispensable for osmotic signalling [32]. In addition, we found that ROP6 controls some of the plant response to osmotic stress. Indeed, plant invalidated for *ROP6* has less osmotically-induced lignin deposition in their roots, a limited isotropic growth when grown on hyperosmotic conditions and a modified root elongation in response to stress condition. Thus, we believe that ROP6 is an important factor for plant osmotic signaling, likely acting just after cell osmotic perception, as ROP6 nanodomain formation happens only minutes after cell stimulation.

Upon its activation by osmotic stimulation, we demonstrate that increased ROS accumulation in cells is associated with the formation of a ROP6/RBOHD complex within the PM. Plant expressing GTP-lock form of ROP6 lead to a high ROS accumulation in cells. In this condition, ROP6 nanoclustering and its colocalization with RBOHD happens without any cell stimulationz. These results fit with our FLIM experiment, where RBOH interact preferentially with ROP6 GTP-locked form. On the other hand, mutated ROP6 that are unable to be acylated, lose both the osmotically-induced nanodomain formation and consequently the ROS accumulation after hyperosmotic stimulation. Our results suggest that ROP6 activation itself can initiate the formation of the ROP6/RBOHD complex, but how this could work mechanistically is still an intriguing question. The constitutive active ROP6 (ROP6-CA) was shown by biochemistry to be associated with detergeant resitant membrane together with a slower diffusion [16,18]. This is mediated firstly by the acylation of C23 and C156 residues of the protein with palmitic and/or stearic acids and secondly by the direct binding between lysine residues in ROP6 hypervariable tail and phosphatidylserine (PS) [16,18]. These results suggest that small GTPases have a greater affinity for specific lipid environment when they are activated, which then determine their nanoclustering. Then, because activated ROP6 is interacting with RBOHs, it might drag and/or retain RBOHs protein to ROP6 nanodomains.

Our group has recently described that two ROS machinery are under action in response to osmotic stimulation, which includes the involvement of two isoforms of NADPH oxidase, RBOHD and F^9^. Our results suggest that ROP6 is an upstream regulator for both ROS generating pathways. But, how recruitment of RBOHs in ROP6 nanodomains can regulate ROS accumulation is still unclear. Because of their ability to generate potentially harmful oxygen radical, RBOHs activity is tightly controlled in cells. This is particularly well described for pathogen elicitors, where several kinases including BIK1 and CPK5 are necessary for PTI mediated ROS accumulation and can directly phosphorylate RBOHD N-terminus [33,34]. The change of RBOHs PM localization mediated by ROP6, could then participate in kinases interaction and consequently alter its phosphorylation/dephosphorylation kinetic. Also, RBOHD and F contain EF-hands that can directly bind calcium and are essential for RBOHs activity [35,36]. Within the cell membrane, calcium gradient might exist in the vicinity of membrane transporter [37]. Therefore, recruitment of RBOHs proteins in ROP6-containing nanodomains could alter RBOHs calcium micro-environment a thereby regulating its activity. In addition, RBOHs dimerization was observed from purified OsRBOHB N-terminus but also suggested from step bleaching experiment done *in vivo* [6,38]. Interestingly, we observed an epistatic interaction between *rbohD* and *rbohF* for osmotically induced ROS, suggesting that RBOHD and F might form heteromers [11]. Similar observations were recently described also for ROS triggered upon cell ablation [39]. We speculate that co-clustering of RBOHD and RBOHF in ROP6-containing nanodomains could increase their probability to form functional heteromers.

Rho GTPases are generally seen as the neck of an hourglass for signal integration at the cell surface. Indeed, multiple input pathways converge on a single Rho GTPase, leading to various downstream cellular outputs, which are often specific to the upstream signal. How signaling specificity is achieved in this context is an outstanding unresolved question. In our work, we found that a single ROP isoform could, in response to different stimuli e.g. auxin and osmotic stimulus, generate very similar nanodomains in terms of shape or cellular density. Nevertheless, we also found that these nanoclusters differ in their composition at least for RBOH proteins. Therefore, the segregation of signaling components in distinct plasma membrane nanodomains can generate signal specificity downstream of a single small GTPase. How this discrimination happens still remains an open question. It could be because of specific lipid environment or/and recruitment of additional proteins that will participate in the stabilization of ROP6/RBOH complex.

## MATERIEL AND METHODS

### Growing condition and plant material

Arabidopsis thaliana Col-0 accession was used as the wild-type (WT) reference background throughout this study. Plants were stratified for 2 days at 4°C and grown vertically on agar plates containing half-strength Murashige and Skoog (½ MS) medium supplemented with 1% (w/v) sucrose and 2.5mM MES-KOH pH6 for 5 days at 22°C in a 16-h light/8-h dark cycle with 70% relative humidity and a light intensity of 200μmol·m−2·s−1, prior to use. N.nicotiana used for transient expressing were grown in soil at 22°C in a 8-h light/16-h dark cycle with 70% relative humidity and a light intensity of 200μmol·m−2·s−1. The following lines were published before: rop6.2 [28], rop2.1 [40], rop4.1 [41], rop6.2xrop2.1 [42], rop6.2xrop2.1xROP4RNAi [42], rop6xpROP6:mCit-ROP6 [18], p35S:EOS-ROP6 [18], p35S:EOS-ROP6-CA [18], mpRBOHD:nls-GUS-GFP [19] and pRBOHF:mcherry-RBOHF [19]. For root architecture analyses, seedlings were grown on vertical square 12×12 cm Petri dishes in a self-contained imaging unit equipped with a 16Mpixel linear camera, a telecentric objective and collimated LED backlight. Plants were grown in the imaging automat dedicated growth chamber at 23°C in a 16-h light/8-h dark cycle with 70% relative humidity and a light intensity of 185 μmol·m−2·s−1 (Vegeled Floodlight, Colasse Seraing – Belgium). Plates were imaged every four hours allowing fine kinetic analysis.

### Cloning and plant transformation

The vector ROP6g/pDONRP2RP3, which includes the full ROP6 genomic sequence from ATG to the end of its 3’UTR (ROP6g – At4g35020) [18] was amplified with the following overlapping to generate either ROP6gCA/pDONRP2R (G15V) or ROP6gDN/DONRP2RP3 (T20N). ROP6gCA/pDONRP2R-P3 and ROP6gDN/pDONRP2R-P3 were then recombine by LR multisite reaction with ROP6prom/pDONRP4P1R [18], mCITRINEnoSTOP/pDONR221 [43] and pB7m34GW [44] to generate pROP6:mCit-ROP6-CA and pROP6:mCit-ROP6-DN vectors, respectively. ROP6g/pDONRP2RP3 was amplified with overlapping primers to generate ROP6gC21S-C156S/pDONRP2RP3. ROP6gC21S-C156S/pDONRP2R-P3 was then recombined by LR multisite reaction with 2×35Sprom/pDONRP4P1R [45], mEOS2noSTOP/pDONR221[18] and pB7m34GW [44] to generate p35S:mEOS2-ROP6C21S-C156S. ROP6gC21S-C156S/pDONRP2R-P3 was also recombined by LR multisite reaction with ROP6prom/pDONRP4P1R [18], mCITRINEnoSTOP/pDONR221 [43] and pB7m34GW [44] to generate ROP6prom:mCITRINE-ROP6C21S-C156S. The coding sequence of RBOHD (At5g47910), RBOHF (At1g64060), ROP6 (At4g35020), ROP6-CA (G15V) and ROP6-DN (T20N) were PCR amplified and inserted into pENTR/D-TOPO. pB7WGF2 and pB7WGR2vector were used as destination vector for respectively GFP and RFP fusion. The different binary were use either for transient expression in tobacco [46] or to generate stable Arabidopsis plants by floral dip method either in col(0) or *rop6.2* [47].

### Osmotic and Pharmacological Treatments

Plantlets were bathed in a liquid MS/2 medium for 30 min to allow recovery from transplanting. When indicated, pre-treatment with DPI (30min, 20µM), BPDS (50 µM, 30min) or NAA (10µM, 1 hour) was added to the media. Then, plantlets were gently transferred for an additional 15 min with 5 μM of ROS dye dehydroethidium (DHE), with or without the corresponding inhibitors, into MS/2 medium (0 MPa), MS/2 medium plus 100 mM sorbitol (−0.26 MPa) for mild stress or MS/2 medium plus 300 mM sorbitol (−0.75 MPa) for severe osmotic stress.

### Western blot

Tissue from 5days old Col0, rop6.2xmCit-ROP6 and GFP-ROP6 plantlets was grinded with liquid nitrogen to a fine powder and resuspended in 1 mL/g powder of RIPA extraction buffer (150 mM NaCL, 50mM Tris-HCl, pH-8, 0.1% SDS, 0.5% Na deoxycholate, 1% Triton x-100, 2mM leupeptin, 1mM PMSF and 5mM DTT). Western blot analysis was performed with antibodies diluted in blocking solution (1% BSA in 0.1% Tween-20 and PBS) at the following dilutions: α-GFP-HRP 1:2000. Whole protein quantity was revealed with Commasie blue stain.

### Sample clarification and phloroglucinol staining

Seedlings, vertically grown in half-strength MS-agar plates for 5 days were transfer on control (MS/2) or 300mM sorbitol plates for 24 hours. Plantlets were treated accordingly to Malamy et al., 1997 [48]. In brief, they were incubated in 0.24 M of HCl prepared in 20% ethanol, at 80°C for 15 minutes, and then transferred in a solution 7% NaOH in 60% ethanol for another 15 minutes at room temperature. The incubated seedlings are rehydrated in subsequent baths for 5 minutes in 40%, 20% and 10% ethanol and infiltrated after in 5% ethanol/25% glycerol for 15 minutes. Alternatively, root samples were stained with phloroglucinol as in Prajakta Mitra et al., 2014 [20].

### ROS and autofluorescence quantification

Observations were performed on the root elongation zone using an Axiovert 200M inverted fluorescence microscope (20×/0.5 objective; Zeiss), with 512/25-nm excitation and 600/50 emission filters for DHE staining and with 475/28 nm excitation and 530/25 nm emission for lignin stained samples. Exposure time was 500 ms. Images were acquired using a CCD camera (Cooled SNAP HQ; PhotoMetrics), controlled by imaging software (MetaFluor; Molecular Devices). To quantify the intensity of the fluorescence signal, the images were analyzed using ImageJ software. After subtraction of the background noise, an average mean grey value was calculated from epidermal and cortical cells.

### Confocal laser scanning microscopy

Signal from rop6.2xmCit-ROP6, rop6.2xmCit-ROP6 CA and rop6.2xmCit-ROP6 DN was imaged using Leica SP8 microscope with a 40×/1.1 water objective and the 488-nm line of its argon laser was used for live-cell imaging. Fluorescence emission was collected from 500–540 nm for GFP and from 600–650 nm by sequential acquisition when sample where stained 10 min with 2 μM of FM4-64.

### TIRF microscopy

For cluster density analysis, Total Internal Reflection Fluorescence (TIRF) Microscopy was done using the inverted Zeiss microscope and a 100x/1.45 oil immersion. Images were acquired with 50ms exposure time at 50 gain, with 475 nm excitation and 530/25 nm emission. Acquisitions were recorded for 0.5 seconds. Images were Z stacked by average intensity and object detection of GFP-ROP6, GFP-ROP6CA, RbohD-GFP and RbohF-GFP was made using machine learning-based segmentation with Elastik [49]. For colocalization study, TIRF microscopy was done using an inverted Nikon microscope (Eclipse) equipped with azimuthal-TIRFiLas2 system (Roper Scientific) and a 100x/1.49 oil immersion. One hundred images were acquired with 100ms exposure time using sequentially 488nm laser illumination with 425/20 emission filters and 561nm laser with 600/25.

### FRET-FLIM

FRET-FLIM measurements were effectuated by multiphoton confocal microscopy (ZEISS LSM 780) with the method of measuring the lifetime of photons (TCSPC: Time correlated single photon counting) and under a 40x/1.3 oil immersion objective (Peter and Ameer-Beg, 2004). The GFP (donor GFP-RBOHD or GFP-RBOHF) was excited with 920 nm by a pulsating infra-red laser Ti:Saphir (Chameleon ULTRA II, COHERENT) during 90 seconds and the emitted fluorescence was collected by HPM-100 Hybrid detector. The decreasing fluorescence curve was obtained with the SPCImage (Becker-HIckl) software for each zone of interest. The lifetime of the GFP was estimated based on a curve regression, either mono-exponential when the donor was expressed alone and bi-exponential when the donor was expressed together in presence of the acceptor proteins (RFP-ROP6-CA and RFP-ROP6-DN). 3 biological repetitions were done and for every biological replicate, 5 cells were analyzed.

### sptPALM

Root cells were observed with a homemade total internal reflection fluorescence microscope equipped with an electron-multiplying charge-coupled device camera (Andor iXON XU_897) and a 100×/1.45 oil immersion objective. The coverslips (Marienfeld 1.5H) were washed sequentially with 100% ethanol, acetone and water. Then, they were bathed with a 1M KOH solution and then ultra-sonified for 30 min. After several wash-outs with MilliQ water, they were dried under Bunsen burner flame. The laser angle was adjusted so that the generation of the evanescence waves give a maximum signal-to-noise ratio. The activation of the photoconvertible tagged mEOS2-ROP6, mEOS2-ROP6-CA and mEOS2-ROP6^C21A/C156A^ was done by a low-intensity illumination at 405 nm (OBIS LX 50mW; Coherent), 561 nm (SAPPHIRE 100mW; Coherent) emission combine a with 600/50 (Chroma) emission filter was used for image acquisition [11]. Ten-thousand images were recorded per region of interest and streamed into LabVIEW software (National Instruments) at 20ms exposure time. 10 to 20 cells/ treatment were analysed out of three biological replicates.

### Single-Particle Tracking and Vonoroi Tessellation

Individual single molecules were localized and tracked using the software MTT [50]. Dynamic properties of single emitters in root cells were then inferred from the tracks using homemade analysis software written in MatLab (The MathWorks) [11]. From each track, the MSD was computed. To reduce the statistical noise while keeping a sufficiently high number of trajectories per cell, tracks of at least five steps (i.e. ≥ 6 localizations) were used. Missing frames due to mEOS2 blinking were allowed up to a maximum of three consecutive frames. The diffusion coefficient D was then calculated by fitting the MSD curve using the first four points. For the clustering analysis, the positions retourned by MTT of each mEOS2 detection were used as input to the SR-Tesseler software [25]. Correction for multiple detection was made based on recommendation from Levet et al., 2015 [25]. The local densities of each track were calculated as the invert of their minimal surface. Then, nanoclusters size, relative number of ROP6 molecules in nanodomains and density of nanocluster was calculated after defining region of interest (ROI) where the local density was 50 times higher than the average. Only ROI with at least 25 detections were considered.

### Statistical Analysis

For each condition or treatment, 9–12 cells were analyzed from at least 5–7 different seedlings. All experiments were independently repeated 2–3 times. Data are expressed as mean ± 95% confidence interval. ANOVA followed by Tukey test was done, letters indicate significant differences among means (pvalue<0.001). * p-value below 0.01 Student T-Test. Statistical analyses were performed in GraphPad Prism (GraphPad Software).

## Acknoledgment

We thank the Montpellier Ressources Imagerie and the Histocytology and Plant Cell Imaging Platform for providing the microscope facility.Y.J. was funded by ERC no. 3363360-APPL under FP/2007-2013; Y.J. and A.M. by the innovative project iRhobot from the department of “Biologie et Amélioration des Plantes” (BAP) of INRA.

## Supplemental figure legends

**FigS1: Expression pattern of different *ROP* isoforms and the ROS production phenotype of single and multiple ROP mutants.** (A) Gene expression clustering of the different ROP isoforms based on eFP-browser databases. Green square shows the three isoform highly express in root tissue (*ROP2, ROP4* and *ROP6*). (B) Quantification of ROS accumulation (DHE staining) in control or after 15 minutes of -0.75 MPa treatment in the indicated genotype. Error bars correspond to a confidence interval at 95%. ANOVA followed by Tukey test was done, letters indicate significant differences among means (p-value<0.001). N =3 independent biological replica.

**FigS2: ROP6 participates to lignin accumulation, cell isotropic growth and root elongation in response to osmotic stimulus.** (A) Cell autofluorescence of *rop6.2* and complemented lines expressing mCit-ROP6 under ROP6 endogenous promotor in control plate or after -0.75 MPa treatment for 24 hours. (B) Phloroglucinol staining, that show pink precipitate when in complex with lignin in control condition or after -0.75 MPa treatment for 24 hours. (C) Cell autofluorescence quantification of Col(0) plant exposed for 24 hours to control, -0.26, -0.5, -0.75 MPa. As comparison, cell autofluorescence was also observed in *rbohDxrbohF* line was exposed to -0.75 MPa and Col(0) treated for 1 hour with 1mM H202 treatment. (D) Cell autofluorescence quantification in *rop6.2* and *rop6.2*xmCit-ROP6 in control or treated plate (−0.75 MPa). (E) 2 days after transfer on - 0.75 MPa plate, root cells present inflated cells (arrow). The arrows are located at the point where the root tip was at the time of transfer. (F) Close up view of cells in this zone in control condition or after treatment (−0.75 MPa) for *rop6.2*xmCit-ROP6 or *rop6.2*. (G) Quantification of cell circularity index. (H-N) the complemented line (*rop6.2*xmCit-ROP6) or the mutant *rop6.2* were grown 5 days on control plates and then transferred for 4 more days in either control condition or on plate supplemented with osmoticum to reach -0.75 MPa of water potential. Relative growth of *rop6.2*xmCit-ROP6 or *rop6.2* in control (J) or in -0.75 MPa plate (K). Quantification of the primary root length (L), lateral density (M) and lateral root length (N) of *rop6.2*xmCit-ROP6 or *rop6.2* grown on -0.75 MPa plate. Error bars correspond to a confidence interval at 95%. ANOVA followed by Tukey test was done, letters indicate significant differences among means (pvalue<0.001). N=3 independent replica. Scale bar 20µm for (A) and 2 mm for (E). N=3 independent biological replica

**FigS3: Expression pattern of *rop6.2*xmCit-ROP6 lines along the root.** (A) Arabidopsis control plant counterstain with calcofluor bright to illustrate the different root zone. Root apical meristem (RAM), elongation zone (EZ), differentiation zone (DZ) and mature zone (MS). (B-E) Representative micrograph of the fluorescent signal observed in *rop6.2* lines complemented with mCit-ROP6 under ROP6 endogenous promoter. (E) mCit-ROP6 signal is mostly visible at the cell PM, reveal by FM4-64 staining. Scale bar 20µm.

**FigS4: Characterization of GFP-ROP6 overexpressing line and localization of *rop6.2*xmCit-ROP6, *rop6.2*xmCit-ROP6-CA and *rop6.2*xmCit-ROP6-DN.** (B) Western blot with antibody against GFP on plant protein extract from Col(0), ROP6 complemented line (*rop6.2*xmCit-ROP6) and ROP6 overexpressing line (ROP6). (B) Confocal micrograph showing the localization of wild type ROP6 (mCit-ROP6), ROP6 constitutive active ROP6 (mCit-ROP6-CA) and the dominant negative ROP6 (mCit-ROP6-DN), and its respective fluorescence at the PM and it is respective quantification (C). CBB, Coomassie brilliant blue. N=2 independent biological replica

**FigS5: ROP6 single-molecule imaging and Vonoroï tessellation.** (A) To verify that we are indeed recording single mEOS2-ROP6 molecules, we plot fluorescence intensity of a typical mEOS2-ROP6 sub-diffractive spot along time. The signal intensity observe is not continuous and the OFF state vary in duration between seconds and milliseconds. This blinking behaviour is typical from single-molecule observation. We also quantify the track duration (B). As expected from single molecules, vast majority of the tracks do not last for more than 0.5 seconds. (C) Picture of Vonoroï diagram, where each point/seeds correspond to a mEOS2-ROP6 localization and edges of Vonoroï cells are represented in white. (D) Segmented region of interest (ROI) with a particle local density greater than log (local density)>3 (ROI appear in red). (C) -Close up view of one ROP6 nanodomain where each blue dots represent one mEOS2-ROP6 localization.

**FigS6: ROP6, RBOHD and RBOHF are expressed in root epidermal cells.** Expression pattern of the translational fusion pROP6:mCit-ROP6 (A) and pRBOHF:mCherry-RBOHF (B) and the transcriptional fusion pRBOHD:nls-GFP-GUS (C).

**FigS7: RBOHF interaction with ROP6 and localization in response to osmotic stimulus.** (A) Quantification of GFP-RBOHF fluorescence life time expressed in transient expression in tobacco leaf epidermal cells, either alone, or co-expressed with the dominant negative (RFP-ROP6-DN) or the constitutive active ROP6 (RFP-ROP6-DN). (B) TIRFM micrograph of cell expressing GFP-RBOHF in control or after 2 minutes treatment with -0.75 MPa solution and quantification of spots density (C). Error bars correspond to a confidence interval at 95%. For (A), ANOVA followed by Tukey test was done, letters indicate significant differences among means (p-value<0.001). * p-value below 0.01 T-Test. N=4 N=3 independent biological replica. Scale bar 10µm.

