## Supplemental data for "A plasma membrane nanoplatform ensures signal specificity during osmotic signaling in plants"

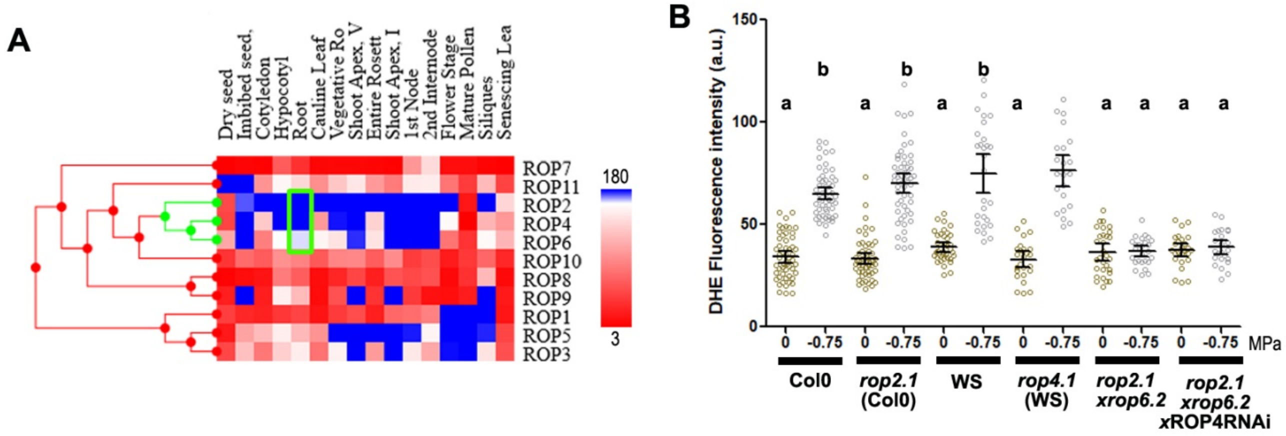


**FigS1: Expression pattern of different *ROP* isoforms and the ROS production phenotype of single and multiple ROP mutants.** (A) Gene expression clustering of the different ROP isoforms based on eFP-browser databases. Green square shows the three isoform highly express in root tissue (*ROP2*, *ROP4* and *ROP6*). (B) Quantification of ROS accumulation (DHE staining) in control or after 15 minutes of -0.75 MPa treatment in the indicated genotype. Error bars correspond to a confidence interval at 95%. ANOVA followed by Tukey test was done, letters indicate significant differences among means (p-value<0.001). N =3 independent biological replica

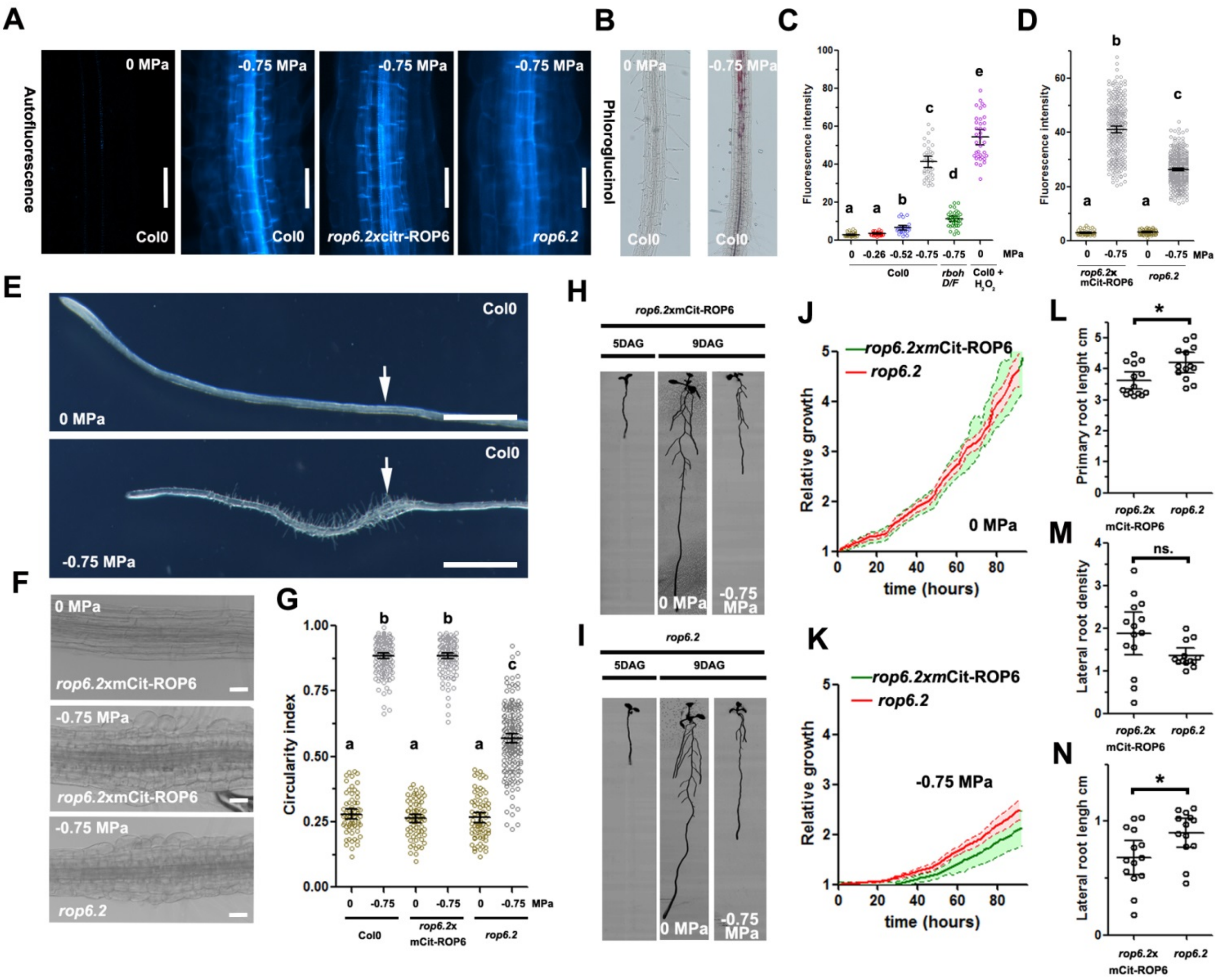

**FigS2: ROP6 participates to lignin accumulation, cell isotropic growth and root elongation in response to osmotic stimulus.** (A) Cell autofluorescence of *rop6.2* and complemented lines expressing mCit-ROP6 under ROP6 endogenous promotor in control plate or after -0.75 MPa treatment for 24 hours. (B) Phloroglucinol staining, that show pink precipitate when in complex with lignin in control condition or after -0.75 MPa treatment for 24 hours. (C) Cell autofluorescence quantification of Col(0) plant exposed for 24 hours to control, -0.26, -0.5, -0.75 MPa. As comparison, cell autofluorescence was also observed in *rbohDxrbohF* line was exposed to -0.75 MPa and Col(0) treated for 1 hour with 1mM H202 treatment. (D) Cell autofluorescence quantification in *rop6.2* and *rop6.2*xmCit-ROP6 in control or treated plate (-0.75 MPa). (E) 2 days after transfer on -0.75 MPa plate, root cells present inflated cells (arrow). The arrows are located at the point where the root tip was at the time of transfer. (F) Close up view of cells in this zone in control condition or after treatment (-0.75 MPa) for *rop6.2*xmCit-ROP6 or *rop6.2*. (G) Quantification of cell circularity index. (H-N) the complemented line (*rop6.2*xmCit-ROP6) or the mutant *rop6.2* were grown 5 days on control plates and then transferred for 4 more days in either control condition or on plate supplemented with osmoticum to reach -0.75 MPa of water potential. Relative growth of *rop6.2*xmCit-ROP6 or *rop6.2* in control (J) or in -0.75 MPa plate (K). Quantification of the primary root length (L), lateral density (M) and lateral root length (N) of *rop6.2*xmCit-ROP6 or *rop6.2* grown on -0.75 MPa plate. Error bars correspond to a confidence interval at 95%. ANOVA followed by Tukey test was done, letters indicate significant differences among means (pvalue<0.001). N=3 independent replica. Scale bar 20µm for (A) and 2 mm for (E). N=3 independent biological replica

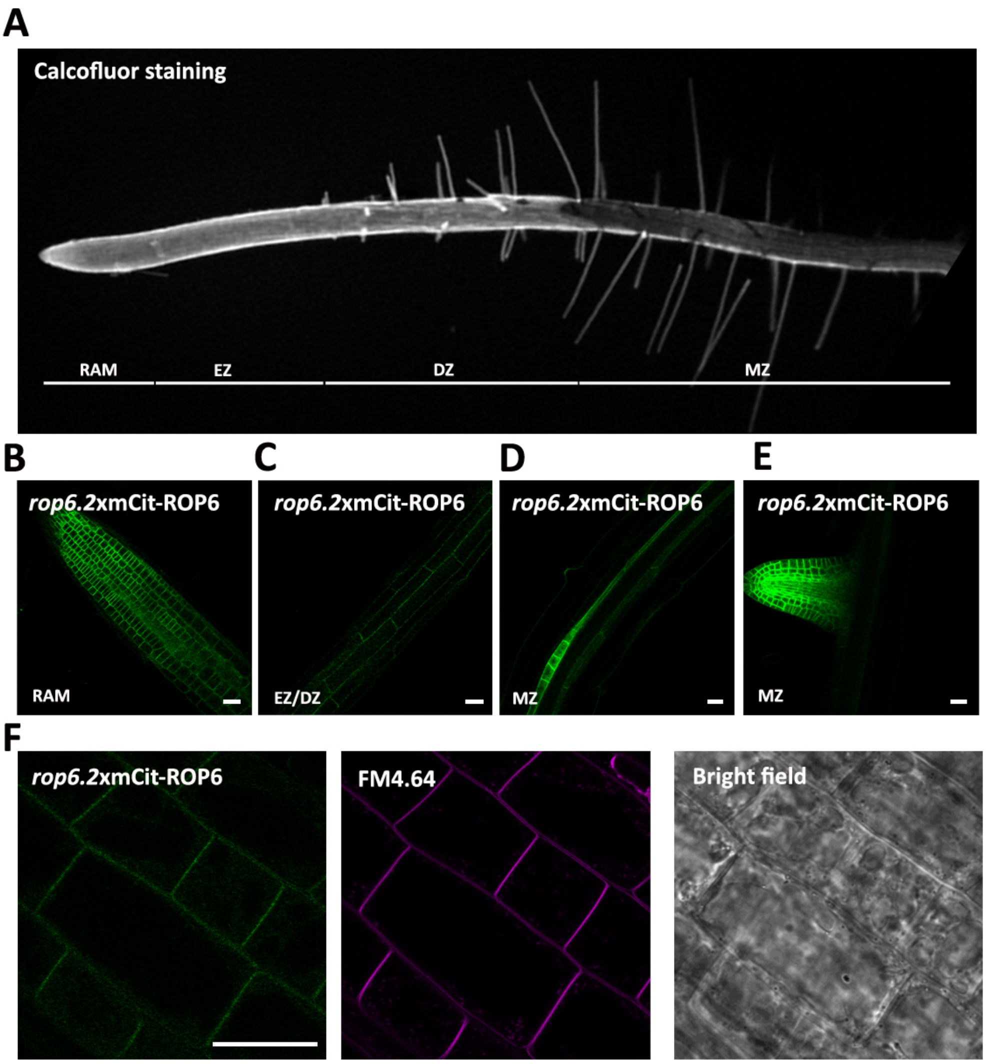


**FigS3: Expression pattern of *rop6.2*xmCit-ROP6 lines along the root.** (A) Arabidopsis control plant counterstain with calcofluor bright to illustrate the different root zone. Root apical meristem (RAM), elongation zone (EZ), differentiation zone (DZ) and mature zone (MS). (B-E) Representative micrograph of the fluorescent signal observed in *rop6.2* lines complemented with mCit-ROP6 under ROP6 endogenous promoter. (E) mCit-ROP6 signal is mostly visible at the cell PM, reveal by FM4-64 staining. Scale bar 20µm.

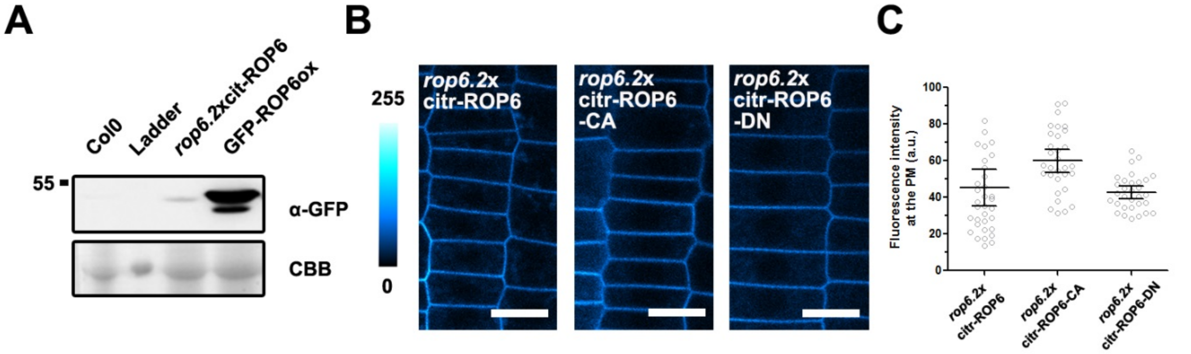


**FigS4: Characterization of GFP-ROP6 overexpressing line and localization of *rop6.2*xmCit-ROP6, *rop6.2*xmCit-ROP6-CA and *rop6.2*xmCit-ROP6-DN.** (B) Western blot with antibody against GFP on plant protein extract from Col(0), ROP6 complemented line (*rop6.2*xmCit-ROP6) and ROP6 overexpressing line (ROP6). (B) Confocal micrograph showing the localization of wild type ROP6 (mCit-ROP6), ROP6 constitutive active ROP6 (mCit-ROP6-CA) and the dominant negative ROP6 (mCit-ROP6-DN), and its respective fluorescence at the PM and it is respective quantification (C). CBB, Coomassie brilliant blue. N=2 independent biological replica

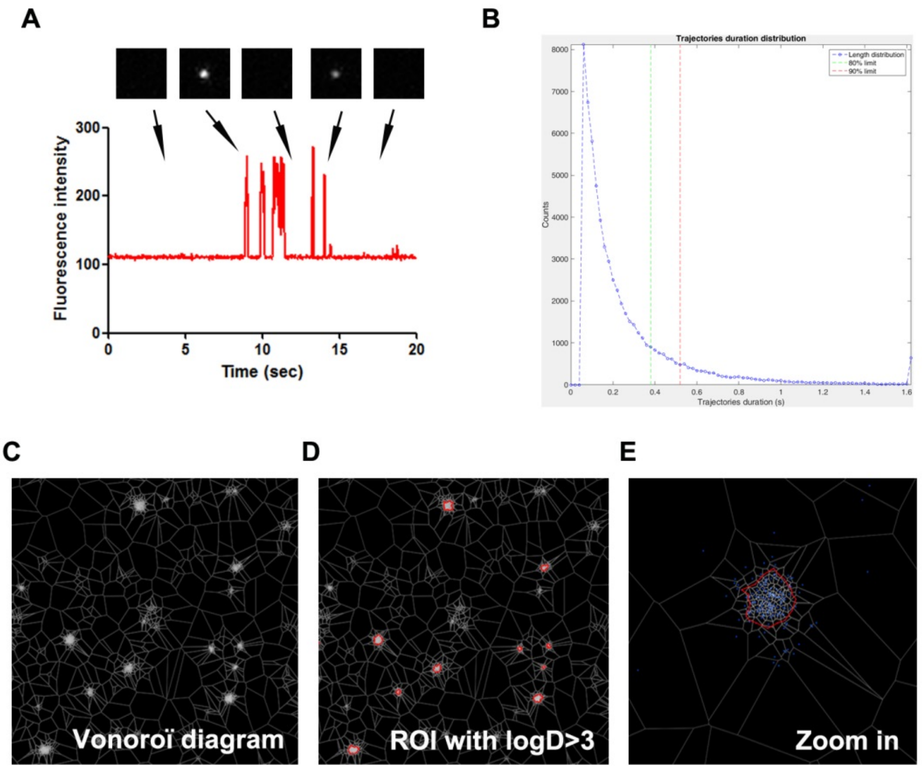


**FigS5: ROP6 single-molecule imaging and Vonoroï tessellation.** (A) To verify that we are indeed recording single mEOS2-ROP6 molecules, we plot fluorescence intensity of a typical mEOS2-ROP6 sub-diffractive spot along time. The signal intensity observe is not continuous and the OFF state vary in duration between seconds and milliseconds. This blinking behaviour is typical from single-molecule observation. We also quantify the track duration (B). As expected from single molecules,  vast majority of the tracks do not last for more than 0.5 seconds. (C) Picture of Vonoroï diagram, where each point/seeds correspond to a mEOS2-ROP6 localization and edges of Vonoroï cells are represented in white. (D) Segmented region of interest (ROI) with a particle local density greater than log (local density)>3 (ROI appear in red). (C) -Close up view of one ROP6 nanodomain where each blue dots represent one mEOS2-ROP6 localization.

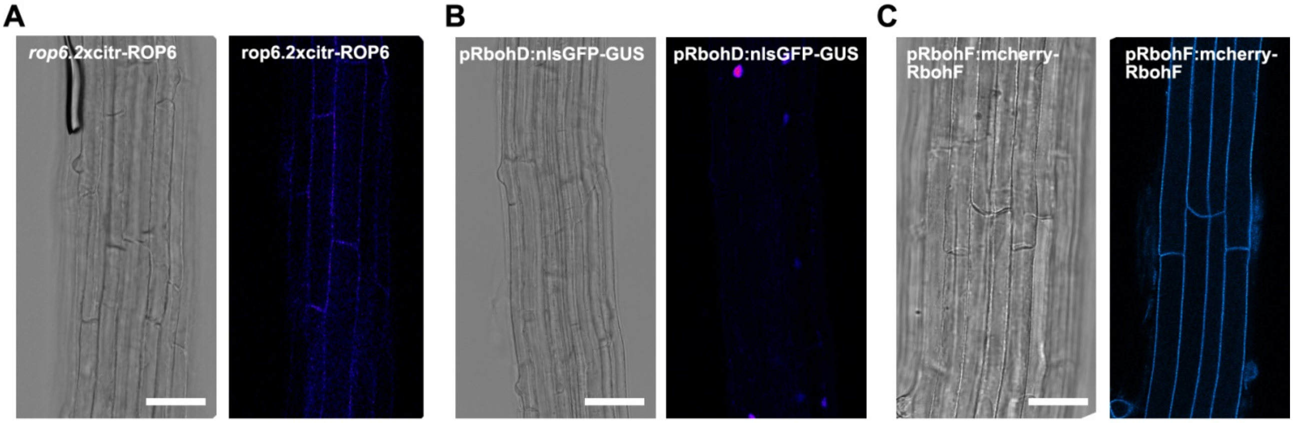


**FigS6: ROP6, RBOHD and RBOHF are expressed in root epidermal cells.** Expression pattern of the translational fusion pROP6:mCit-ROP6 (A) and pRBOHF:mCherry-RBOHF (B) and the transcriptional fusion pRBOHD:nls-GFP-GUS (C).

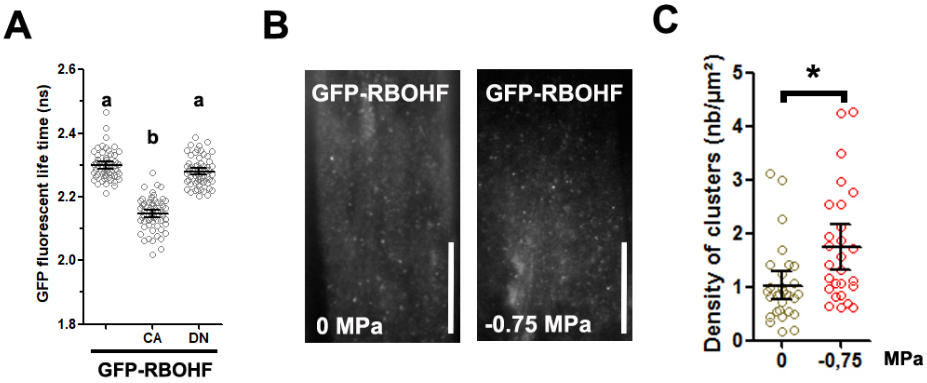


**FigS7: RBOHF interaction with ROP6 and localization in response to osmotic stimulus.** (A) Quantification of GFP-RBOHF fluorescence life time expressed in transient expression in tobacco leaf epidermal cells, either alone, or co-expressed with the dominant negative (RFP-ROP6-DN) or the constitutive active ROP6 (RFP-ROP6-DN). (B) TIRFM micrograph of cell expressing GFP-RBOHF in control or after 2 minutes treatment with -0.75 MPa solution and quantification of spots density (C). Error bars correspond to a confidence interval at 95%. For (A), ANOVA followed by Tukey test was done, letters indicate significant differences among means (p-value<0.001). * p-value below 0.01 T-Test. N=4 N=3 independent biological replica. Scale bar 10µm.
